# Mitochondrial citrate metabolism and efflux regulates trophoblast differentiation

**DOI:** 10.1101/2023.01.22.525071

**Authors:** Renee M. Mahr, Snehalata Jena, Sereen K. Nashif, Alisa B. Nelson, Adam J. Rauckhorst, Ferrol I. Rome, Ryan D. Sheldon, Curtis C. Hughey, Patrycja Puchalska, Micah D. Gearhart, Eric B. Taylor, Peter A. Crawford, Sarah A. Wernimont

**Affiliations:** Department of Obstetrics, Gynecology and Women’s Health, University of Minnesota, Minneapolis MN; Division of Molecular Medicine, Department of Medicine, University of Minnesota, Minneapolis MN; Fraternal Order of Eagles Diabetes Research Center, University of Iowa, Iowa City IA; Department of Molecular Physiology and Biophysics, University of Iowa, Iowa City IA; Department of Biochemistry, University of Iowa, Iowa City IA; Department of Genetics, Cell Biology and Development, University of Minnesota, Minneapolis MN; Department of Biochemistry, Molecular Biology and Biophysics, University of Minnesota, Minneapolis MN

**Keywords:** trophoblast, cytotrophoblast, syncytiotrophoblast, differentiation, mitochondria, mitochondrial citrate transporter, SLC25A1, CIC, citrate, metabolomics, isotope tracing metabolomics, RNA seq, BeWo

## Abstract

Cytotrophoblasts fuse to form and renew syncytiotrophoblasts necessary to maintain placental health throughout gestation. During cytotrophoblast to syncytiotrophoblast differentiation, cells undergo regulated metabolic and transcriptional reprogramming. Mitochondria play a critical role in differentiation events in cellular systems, thus we hypothesized that mitochondrial metabolism played a central role in trophoblast differentiation. In this work, we employed static and stable isotope tracing untargeted metabolomics methods along with gene expression and histone acetylation studies in an established cell culture model of trophoblast differentiation. Trophoblast differentiation was associated with increased abundance of the TCA cycle intermediates citrate and α-ketoglutarate. Citrate was preferentially exported from mitochondria in the undifferentiated state but was retained to a larger extent within mitochondria upon differentiation. Correspondingly, differentiation was associated with decreased expression of the mitochondrial citrate transporter (CIC). CRISPR/Cas9 disruption of the mitochondrial citrate carrier showed that CIC is required for biochemical differentiation of trophoblasts. Loss of CIC resulted in broad alterations in gene expression and histone acetylation. These gene expression changes were partially rescued through acetate supplementation. Taken together, these results highlight a central role for mitochondrial citrate metabolism in orchestrating histone acetylation and gene expression during trophoblast differentiation.

## Introduction

Successful pregnancies require the development of a well-functioning placenta. Critical steps in placental development include the formation of placental villi, where a layer of multi-nucleated syncytiotrophoblasts separate fetal vasculature from maternal blood.^1^ Syncytiotrophoblasts are critical for the health of the pregnancy, transporting nutrients to and from the fetus, facilitating maternal-fetal gas exchange, and synthesizing hormones necessary to sustain the pregnancy.^2,3^ Throughout pregnancy, cytotrophoblasts divide and fuse to maintain this syncytial layer through tightly regulated processes requiring the coordination of signaling, transcriptional and cellular fusion machinery.^4,5^ Abnormalities in syncytialization are associated with pregnancy complications including fetal growth abnormalities, preeclampsia, gestational diabetes, and stillbirth.^6,7^

During the process of trophoblast differentiation, coordinated transcriptional re-programming increases expression of gene products involved in syncytia formation and function including HCG (*CGA* and *CGB*), fusion proteins Syncytin 1 and Syncytin 2 (*ERVW* and *ERVFRD*), and transcription factors including those encoded by *GCM1, OVOL1*, and *GATA2*.^4,5,8-10^ In addition, gene products required for cytotrophoblast lineages including the transcription factors, *TEAD4* and *TP63* are downregulated. Epigenetic modifications, including histone acetylation, play a critical role in this process.^11^

Histone acetylation is a post-translational modification typically associated with increased chromatin accessibility and gene expression. Recent work demonstrates that cytotrophoblasts maintain high levels of histone acetylation at multiple lysine residues, which decrease upon differentiation.^11^ Histone deacetylation is required for trophoblast differentiation and is mediated by histone deacetylases 1 and 2. While histone acetylation is a critical regulator of trophoblast differentiation, how changes in substrate availability impacts this process is unclear.

Mitochondria regulate cellular differentiation events across multiple systems and control entry and exit of metabolites across the inner mitochondrial membrane through regulated networks of mitochondrial transport proteins.^12-16^ This allows mitochondria to direct carbon resources to support diverse cellular functions including energy generation, gene expression, and production of epigenetic co-factors.^13-16^ Recent work using high-resolution respirometers has demonstrated that cytotrophoblasts and syncytiotrophoblasts differ in their cellular metabolism.^17,18^ However, differences in oxygen consumption rates from supplied substrates do not fully capture how mitochondrial routing of metabolites may alter cellular outcomes. Identifying mechanisms by which mitochondria route metabolites is key to understanding how changes in nutrient availability regulate trophoblast differentiation.

In this work, we employ high-resolution liquid chromatography mass spectrometry (LC-MS) to determine how differentiation impacts relative abundance of key metabolites involved in the tricarboxylic acid (TCA) cycle. Additionally, we identify the mitochondrial citrate carrier (CIC) as a key regulator of trophoblast differentiation. Loss of CIC results in broad changes in gene expression and impairments in histone deacetylation. Application of exogenous acetate partially rescues gene expression. Overall, our results demonstrate that trophoblast differentiation is associated with changes in mitochondrial citrate metabolism and efflux, which in turn may regulate gene expression through modulation of histone acetylation.

## Results

### Trophoblast differentiation alters metabolite abundance, energy charge, and glucose utilization

Throughout gestation, cytotrophoblasts fuse to form and renew syncytiotrophoblasts.^4,5,8^ Prior work demonstrates cytotrophoblasts and syncytiotrophoblasts differ in their oxygen consumption rate.^17,18^ However, it is unclear how differentiation impacts relative abundance of key glycolytic and TCA cycle intermediates and energy metabolites. Human BeWo cells have been used to elucidate the molecular mechanisms that regulate syncytia formation. In this model system, both biochemical and morphologic changes canonical to differentiation occur following treatment with the protein kinase A activator, forskolin (FSK) **(Supplemental Figure 1A)**.^19,20^ Biochemical differentiation describes the transcriptional changes and production of hormones such as human chorionic gonadotropin (HCG) key to syncytiotrophoblast function, whereas morphologic differentiation describes cell fusion resulting in syncytia formation. These aspects of trophoblast differentiation can be differentially regulated.^21-24^ Forty-eight hours following treatment with forskolin, BeWo cells increase *CGA, CGB*, and *ERVW* gene expression **(Supplemental Figure 1B)** and undergo morphological changes and increased fusion index consistent with syncytialization **(Supplemental Figure 1C-D)**.

To address how differentiation impacts metabolite abundance upon differentiation, we applied a high-resolution LC-MS approach to quantify relative abundance of glycolytic and TCA intermediates in our BeWo model. Differentiation upon forskolin treatment significantly increased lactate (53%, *p*<0.01), citrate (39%, *p*<0.0001), aconitate (110%, *p*<0.0001), α-ketoglutarate (151%, *p*<0.0001), succinate (70%, *p*<0.0001) and acetyl CoA (53%, *p*<0.01) total pools relative to control cells, while fumarate and malate remained unchanged (**Figure 1A**). To further determine how differentiation impacts energy status of the cell, we quantified AMP, ADP, and ATP levels. Differentiation increased ATP levels by 35% (*p*<0.01) and ADP levels by 60% (*p*<0.05) **(Figure 1B)** indicating increased energy charge.

**Figure 1:**
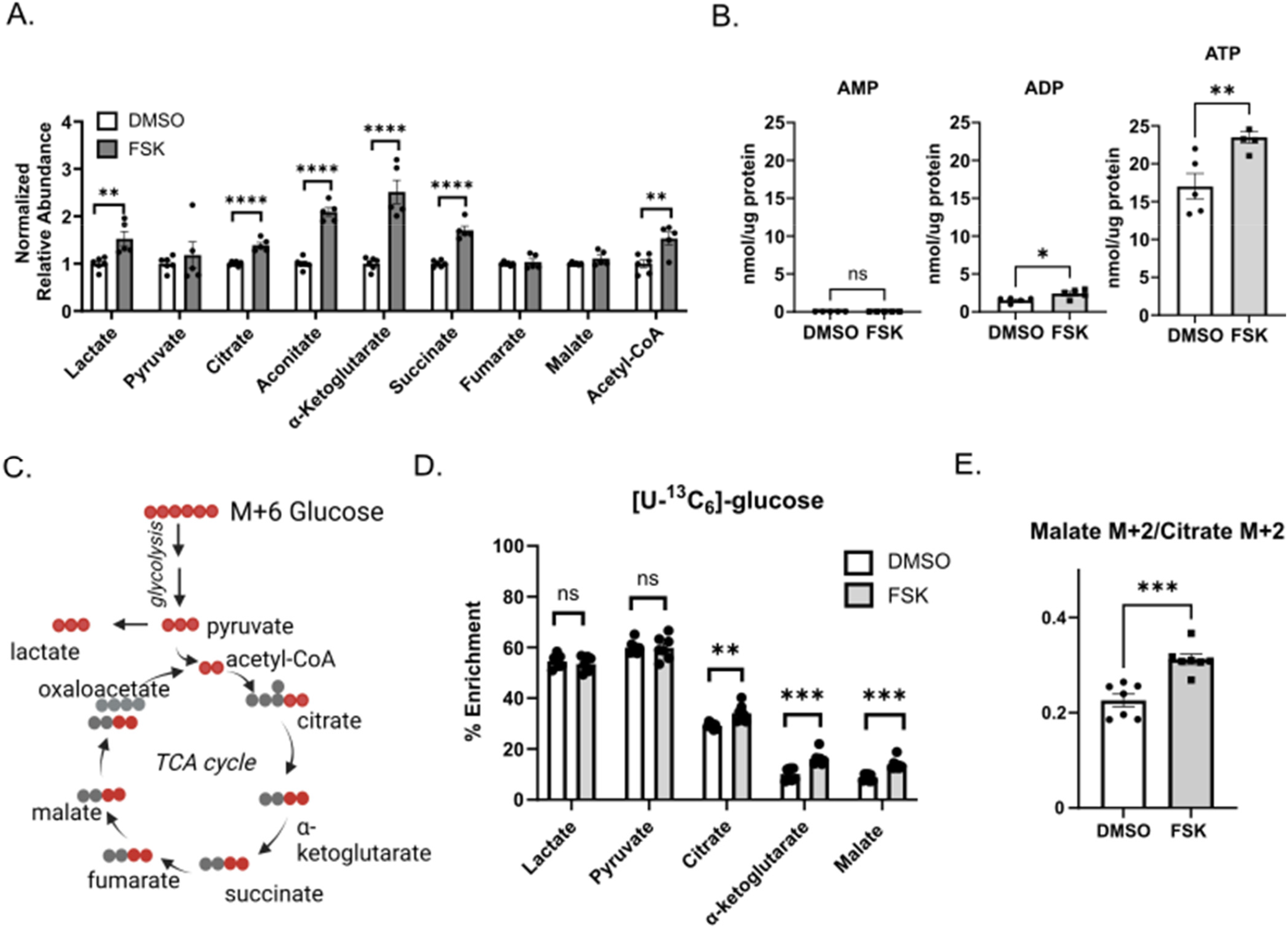
Trophoblast differentiation alters metabolite abundance, energy charge and glucose utilization. A) Relative abundance of citric acid cycle intermediates 48 hours following treatment of BeWo cells with vehicle (0.4% DMSO) or forskolin (FSK, 40 µM). Normalized relative abundance was determined by dividing total signal intensity by mg of total protein and normalizing to DMSO-treated cells. n=5 biological replicates. B) AMP/ADP/ATP levels in nmol/µg protein 48 hours following treatment of BeWo cells with DMSO or 40 uM forskolin (FSK). n=5 biological replicates. C) Graphic depicting path of carbon atoms from glucose after entering the TCA cycle. D) Percent enrichment of [U-^13^C_6_]-glucose in citric acid cycle intermediates 48 hours following treatment of BeWo cells with DMSO or forskolin (FSK). n=6 biological replicates. E) Ratio of Malate M+2 to Citrate M+2 isotopologues, 48 hours following treatment of BeWo cells with DMSO or forskolin (FSK). n=6 biological replicates. All data are representative of mean +/- SEM. *, *p*<0.05; **, *p*<0.01; ***, *p*<0.001; and ****, *p*<0.0001.

Given these initial observations of increased relative abundance of TCA cycle intermediates with differentiation, we investigated how differentiation impacts glucose utilization. Forty-eight hours following treatment of BeWo cells with DMSO or forskolin, we incubated with 17 mM uniformly ^13^C-labeled glucose ([U-^13^C_6_]-glucose) for 6 hours and determined ^13^C-enrichment relative to total pool size using isotope tracing untargeted metabolomics approach **(Figure 1D)**. Both cytotrophoblasts and syncytiotrophoblasts rapidly metabolized glucose to pyruvate and lactate **(Figure 1D)** with approximately 50% of the total pool ^13^C-enriched. Neither lactate nor pyruvate demonstrated forskolin-dependent changes in enrichment. However, approximately 30% of the cytotrophoblast total citrate pool was ^13^C-enriched and this enrichment increased to approximately 35% following forskolin-induced differentiation (*p* < 0.01) **(Figure 1D)**. Glucose derived ^13^C-enrichment into α-ketoglutarate and malate were lower than that of citrate, labeling 5% of the total pool size in cytotrophoblasts and 10% in syncytiotrophoblasts (*p*<0.001). These differentiation-dependent changes in citrate, α-ketoglutarate, and malate ^13^C-enrichment corresponded to the observed increases in total pool size **(Figure 1A)** and suggest that TCA cycle remodeling accompanies differentiation.

To further examine the fate of glucose derived carbon during differentiation, we determined the ratio of the M+2 isotopologue of malate to the M+2 isotopologue of citrate, since this ratio depicts changes in the flux of glucose derived carbon through the TCA cycle **(Figure 1C)**. Trophoblast differentiation significantly increased this ratio suggesting increased forward flux of glucose into the second span of the TCA cycle **(Figure 1E)**. These findings suggest that undifferentiated cytotrophoblasts export mitochondrial citrate to the cytoplasm, whereas differentiated syncytiotrophoblasts retain relatively more citrate to support forward flux through the TCA cycle. These results may reflect activity of the citrate-malate shuttle via a “non-canonical” TCA cycle recently described in embryonic stem cells and myocytes.^25,26^

### Trophoblast differentiation decreases expression of the mitochondrial citrate carrier

Import and export of metabolites through the inner mitochondrial membrane require mitochondrial transport proteins.^12^ Specifically, the mitochondrial citrate carrier (CIC) is responsible for transporting citrate across the inner mitochondrial matrix into the cytoplasm in exchange for malate.^27,28^ Following export of citrate across the inner mitochondrial matrix, it is converted to cytoplasmic acetyl-CoA through the action of ATP citrate lyase (ACLY). Alternatively, cytoplasmic acetyl-CoA can be produced from acetate via ACSS2 **(Figure 2A)**.^29,30^

**Figure 2:**
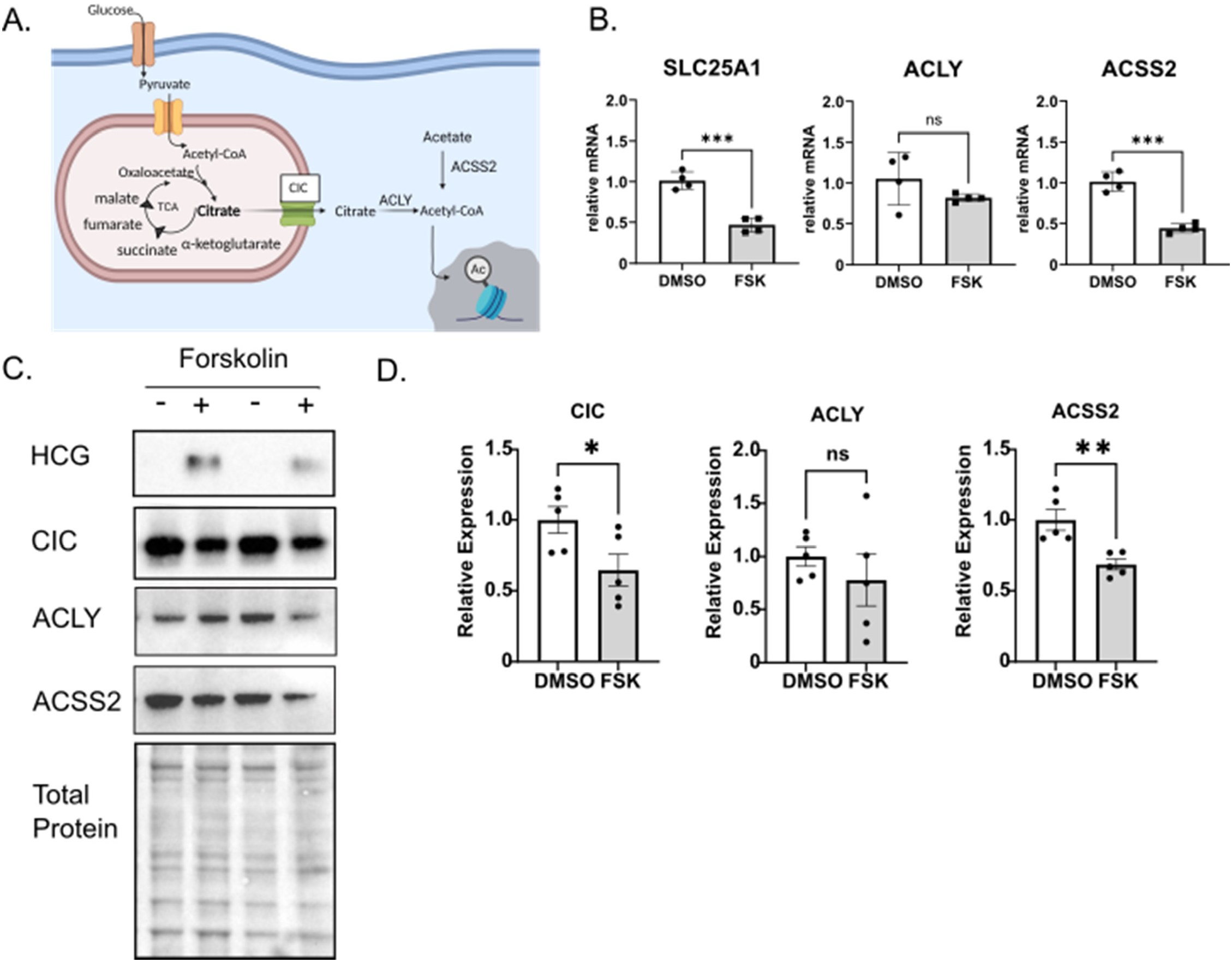
Trophoblast differentiation decreases expression of the mitochondrial citrate carrier. A) Schematic of glucose metabolism to acetyl-CoA. Pyruvate is transported via the mitochondrial pyruvate carrier into the inner mitochondrial membrane where it is converted to acetyl-CoA, fueling the citric acid cycle (TCA). Citrate is transported by the citrate carrier (CIC) to the cytoplasm where ATP citrate lyase (ACLY) converts it to acetyl-CoA, which can be used for histone acetylation. Acetate may contribute to Acetyl-CoA pools through the activity of Acetyl-CoA Synthetase 2 (ACSS2). Figure generated with Biorender. B) qPCR analysis showing relative expression *of SLC25A1, ACLY* and *ACSS2* mRNA following 48 hours treatment of BeWo cells with DMSO or 40 µM Forskolin (FSK). n= 4 biological replicates C) Representative western blot images of HCG, CIC, ACLY, ACSS2 proteins and total protein expression in BeWo cells treated with DMSO or 40 uM forskolin. D) Quantification of *CIC, ACLY*, and *ACSS2* gene expression by qPCR in BeWo cells treated with DMSO or 40 µM Forskolin (FSK). n=5 biologic replicates. Data are representative of mean +/- SEM. *, *p*<0.05; **, *p*<0.01; ***, *p*<0.001; and ****, *p*<0.0001.

Given that mitochondrial citrate efflux may be decreased with differentiation, we next quantified expression of CIC (encoded by *SLC25A1*) during differentiation at mRNA and protein levels. CIC and ACSS2 mRNA and protein levels both decreased following treatment with forskolin **(Figure 2B-D)**. A decrease was not observed for ACLY. Similarly, *SLC25A1* mRNA expression decreased in a trophoblast stem cell model following differentiation indicating this difference in expression is not unique to BeWo cells **(Supplemental Figure 2)**.^31^ Overall, these results suggest that CIC regulates citrate export from mitochondria and partitions citrate and acetyl-CoA during differentiation.

### Loss of mitochondrial citrate carrier impairs trophoblast differentiation and transcriptional reprogramming associated with differentiation

Prior work demonstrates that loss of citrate efflux impairs embryonic stem cell differentiation and that CIC is essential for early trophoblast function.^25,32^ Given decreased citrate export and decreased CIC expression with differentiation, we next determined how loss of mitochondrial citrate carrier prior to differentiation impacts subsequent trophoblast differentiation. Using a CRISPR-Cas9 system,^33^ transduction of three distinct guides targeting *SLC25A1* resulted in complete loss of CIC protein expression compared to empty vector and scrambled non-targeting guide RNA (gRNA)-treated controls **(Figure 3A)**. We subsequently report results from two control and two knockout lines for most assays. To test the impact of CIC on biochemical differentiation, HCG production and gene expression was assessed. HCG following differentiation was 50% lower in CIC KO cells compared to control cells **(Figure 3B)**. Loss of CIC decreased mRNA expression of *CGA, CGB, ERVW* and *ERVFRD* by approximately 60% upon differentiation **(Figure 3C)**. To test the impact of CIC on morphologic syncytialization, BeWo cells were plated on coverslips and stained for E-cadherin and HCG following syncytialization. Loss of CIC did not impair morphologic syncytialization **(Supplemental Figure 3)**, suggesting that CIC uncouples biochemical and morphologic differentiation.

**Figure 3:**
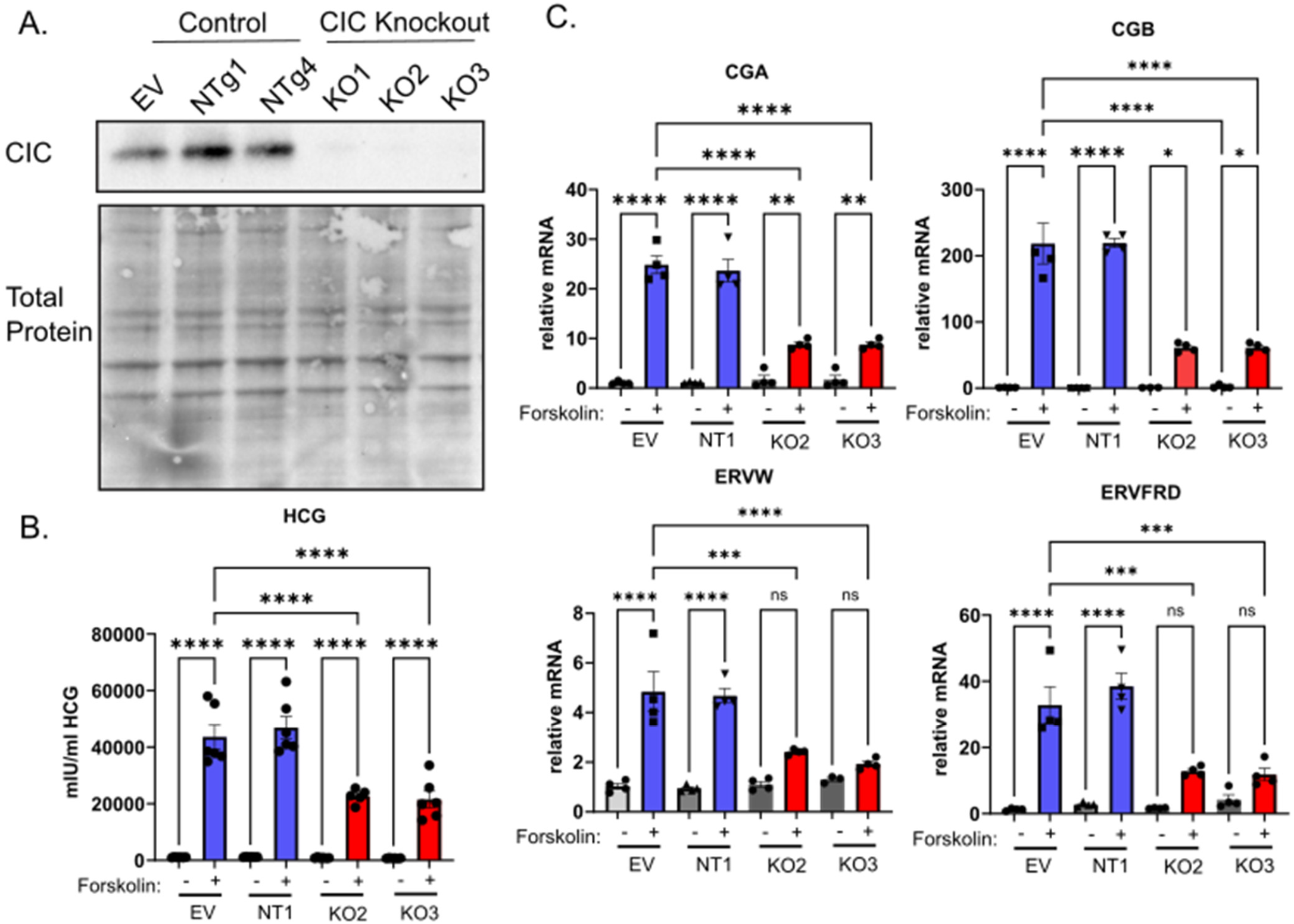
Loss of CIC impairs biochemical differentiation. A) Representative western blot image of CIC and total protein expression in empty-vector, non-targeting, and CIC knockout BeWo cells treated cells. B) ELISA of HCG production in empty-vector, non-targeting, and CIC knockout BeWo cells treated with DMSO or 40 µM Forskolin (FSK). n=3 biological replicates. C) qPCR analysis of *CGA, CGB, ERVW*, and *ERVFRD* gene expression in empty vector, non-targeting, and CIC knockout BeWo cells treated with DMSO or 40 µM Forskolin (FSK). n=3 biological replicates. Data are representative of mean +/- SEM. *, *p*<0.05; **, *p*<0.01; ***, *p*<0.001; and ****, *p*<0.0001.

Given these initial observations highlighting impairments in biochemical differentiation, RNA sequencing was performed following treatment of control or CIC knockout BeWo cells with forskolin or DMSO vehicle. As expected, treatment with forskolin caused dramatic changes in mRNA abundances accounting for 68% of the variance represented by the first principal component (PC1), while loss of CIC is represented within the second principal component (PC2) accounting for 11% of the variance **(Figure 4A)**. Vehicle-treated control and CIC knock-out cells have similar expression along PC1 whereas treatment with forskolin caused a larger shift for control cells compared to the knock-out cells along PC1, consistent with the observations above suggesting that loss of CIC compromises differentiation. Next we used differential expression analysis to identify genes affected by CIC loss. We found 24 genes differentially expressed due to CIC loss in vehicle-treated cells and 88 genes differentially expressed due to CIC loss in forskolin-treated cells using absolute fold change of 1.5, *p* adjusted <0.01. (**Figure 4B-C)**. Using pathway analysis of genes differentially expressed upon treatment with forskolin, loss of CIC decreased expression in pathways that regulate metabolic processes, reproductive system development, and female pregnancy and increased expression in pathways that regulate cytoskeletal reorganization **(Figure 4D)**. Further assessment of these pathways indicated that many observed differences were driven by genes involved in differentiation. Specifically, loss of CIC impaired up-regulation of *CGA, CGB, HSD11B2, ERVW, and ERVFRD* and impaired down-regulation of genes maintaining cytotrophoblasts including *TP63* and *ITGA6* **(Figure 4E)**. Alterations in expression of these genes in control and CIC KO were selectively confirmed using qPCR **(Figure 4F)**.

**Figure 4:**
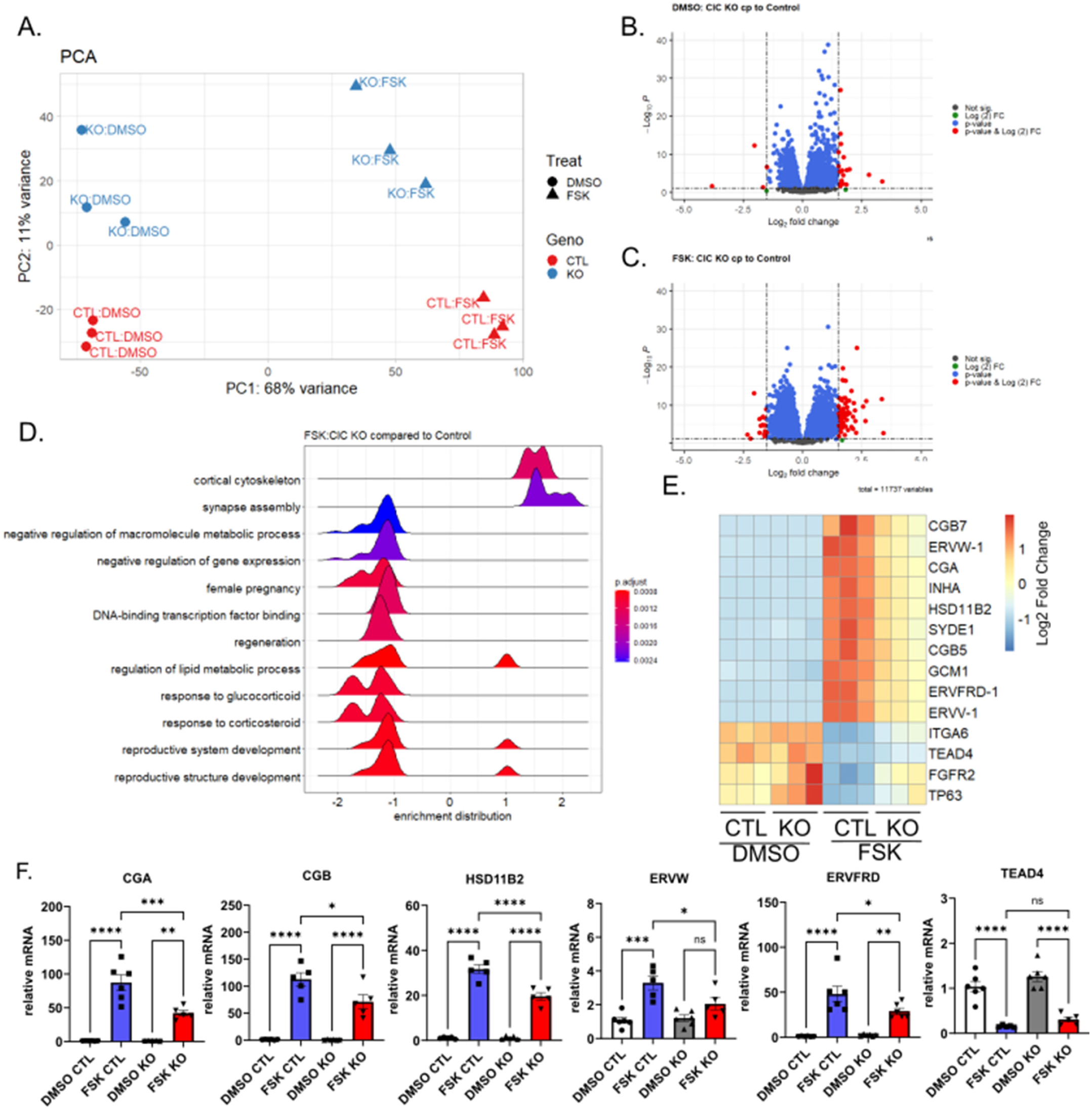
Loss of CIC alters gene expression following trophoblast differentiation. A) PCA plot comparing control (red) and CIC knockout BeWo cells (blue) treated with DMSO (circle) or 40 µM forskolin (FSK) (triangle). B) Volcano plot comparing control and CIC knockout BeWo cells treated with DMSO. C) Volcano plot comparing control and CIC knockout BeWo cells treated with 40 µM forskolin (FSK). D) Pathway analysis comparing control and CIC knockout BeWo cells treated with 40 µM forskolin (FSK). E) Heat map of relative gene expression changes of DMSO and 40 µM forskolin treated control and CIC knockout BeWo cells. F) *CGA, CGB, HSD11B2, ERVW, ERVFRD*, and *TEAD4* gene expression by qPCR in control and CIC knockout BeWo cells treated with DMSO or 40 µM Forskolin (FSK). n=6; Data are representative of mean +/-SEM. *, *p*<0.05; **, *p*<0.01; ***, *p*<0.001; and ****, *p*<0.0001.

### Loss of CIC dysregulates histone acetylation

Prior work demonstrates that histone deacetylation is required for trophoblast differentiation, with global decreases in histone acetylation observed following differentiation in primary trophoblasts, trophoblast stem cells, and BeWo cells.^30^ Using ChIP sequencing, decreased histone acetylation was observed at genes traditionally associated with cytotrophoblastic state and increased histone acetylation observed at genes associated with differentiated states.^11^ While histone deacetylases 1 and 2 regulate these deacetylation events, how changes in acetyl-CoA substrate impact this process in trophoblasts remains unknown. Given the broad changes in gene expression observed with loss of the mitochondrial citrate carrier, we assessed CIC’s impact on histone acetylation during trophoblast differentiation. Similar to prior results, differentiation decreased global histone acetylation and acetylation at lysine 9 in control cells **(Figure 5A-B)**. Strikingly, loss of CIC impaired total histone H3 and lysine 9 deacetylation following forskolin treatment **(Figure 5A-B)**. No statistically significant difference in H3 lysine 27 acetylation in control or CIC knockout cells was detected with differentiation. Combined, this demonstrates that CIC regulates histone acetylation during differentiation and abnormalities in histone acetylation may contribute to the dysregulation of gene expression following loss of CIC expression.

**Figure 5:**
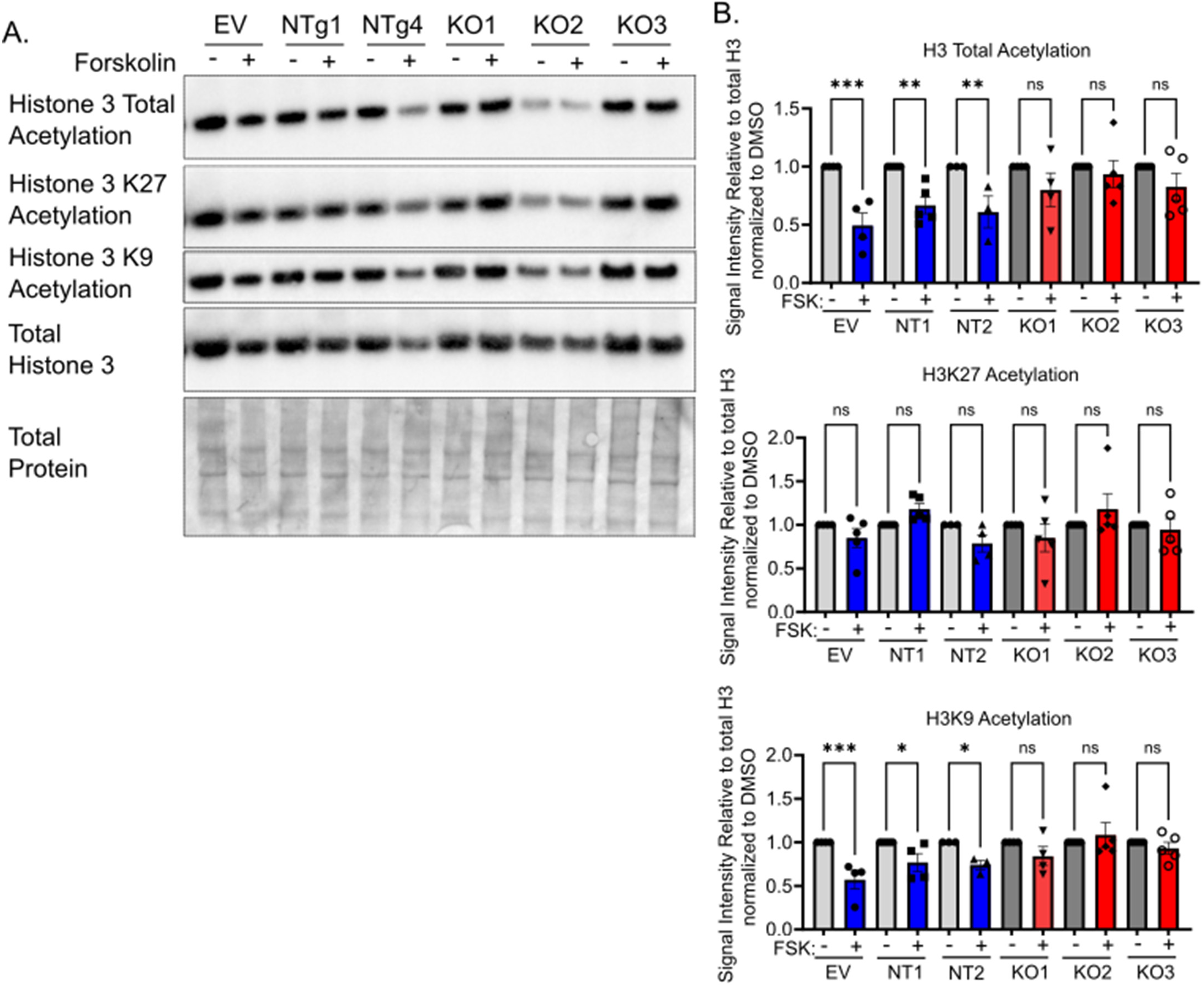
Impaired Histone 3 deacetylation following differentiation of CIC knockout BeWo cells with forskolin. A) Representative western blot of Histone 3 total acetylation, Histone 3 lysine 9 acetylation (H3AcK9), Histone 3 lysine 27 acetylation (H3AcK27), total histone 3 and total protein expression in BeWo cells treated with DMSO or 40 µM forskolin. B) Quantification of forskolin to DMSO relative intensity of histone acetylation to total histone 3 acetylation. n= 5, Data are representative of mean +/-SEM. *, *p*<0.05; **, *p*<0.01; ***, *p*<0.001; and ****, *p*<0.0001.

### Acetate partially rescues gene expression following trophoblast differentiation

Citrate export across the inner mitochondrial membrane contributes to the acetyl-CoA pool required for histone acetylation through the actions of ATP-citrate lyase (encoded by *ACLY*). Acetate may also be converted to acetyl-CoA through ACSS2 in the cytoplasm to support histone acetylation. We assessed whether acetate could rescue the dysregulated differentiation observed in CIC knockout cells. To do this, control and CIC knockout cell lines were treated with 1 mM acetate and the expression of key differentiation markers was quantified via qPCR. While acetate rescued expression of *CGA, CGB* and *ERVW*, it did not rescue expression of *HSD11B2* or *ERVFRD* **(Figure 6)**. This partial rescue suggests that loss of mitochondrial citrate export may have locus-specific effects and that cytoplasmic acetyl-CoA may be a critical mediator of impaired differentiation following loss of CIC.

**Figure 6:**
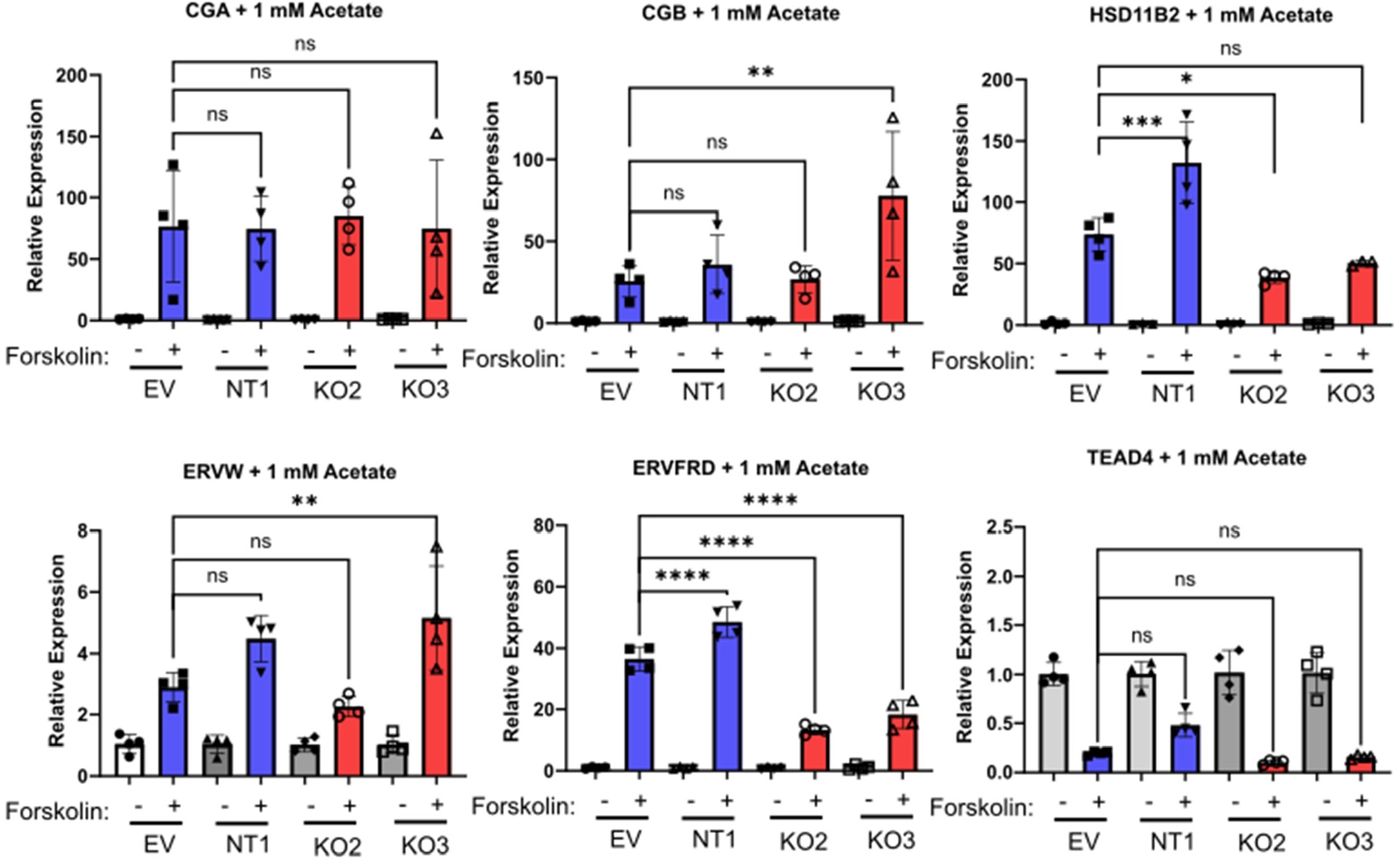
Acetate partially rescues markers of differentiation in CIC knockout cytotrophoblasts. *CGA, CGB, HSD11B2, ERVW, ERVFRD*, and *TEAD4* gene expression by qPCR in empty-vector, non-targeting, and CIC knockout BeWo cells treated with DMSO or 40 µM Forskolin (FSK), in the presence of 1 mM acetate. n=4 biologic replicates. Data are representative of mean +/- SEM. *, *p*<0.05; **, *p*<0.01; ***, *p*<0.001; and ****, *p*<0.0001.

### Loss of mitochondrial citrate efflux results in impaired metabolic programming of differentiating cytotrophoblasts

Given that addition of acetate did not fully rescue differentiation in CIC knockout cells, we next assessed how loss of CIC impacts TCA cycle metabolites following differentiation using high-resolution LC-MS. Assessment of static total cellular pools of metabolites showed that loss of CIC decreased citrate in both vehicle- and forskolin-treated cells **(Figure 7A)**. However, no CIC-dependent differences were observed in α-ketoglutarate, fumarate or malate abundance compared to empty vector or scrambled gRNA-treated control cells. Surprisingly, total cellular acetyl-CoA pool sizes were preserved in CIC knockout cells compared to scrambled gRNA-treated control cells, and increased with forskolin treatment, which may reflect increased mitochondrial localization of acetyl-CoA. To test this, isotope tracing untargeted metabolomics with [U-^13^C_6_]glucose was performed to determine how loss of citrate efflux impacts incorporation of ^13^C-label into TCA cycle metabolites. Both control and CIC knockout cells readily incorporated glucose-derived carbon into pyruvate to similar levels across all conditions **(Figure 7B)**. With differentiation, ^13^C-enrichment of citrate increased in control cells **(Figure 7C)** consistent with results shown in **Figure 1E**. However, the forskolin-induced increased citrate enrichment observed in control-gRNA cells was not observed in CIC knockout cells following differentiation and the total ^13^C-enrichment was in fact lower than forskolin-treated, empty vector or scrambled gRNA-treated control cells. This decreased enrichment correlated with the smaller citrate pool size observed with CIC knockout **(Figure 7A)**. Additionally, while differentiation increased malate enrichment in both control and CIC knockout cells, lower total enrichment was found following forskolin treatment of CIC knockout cells **(Figure 7D)**. Given that loss of CIC does not diminish total malate pool size, this decrease in malate ^13^C-enrichment likely reflects metabolic compensation from anaplerotic sources other than glucose-sourced pyruvate, e.g., amino acids.

**Figure 7:**
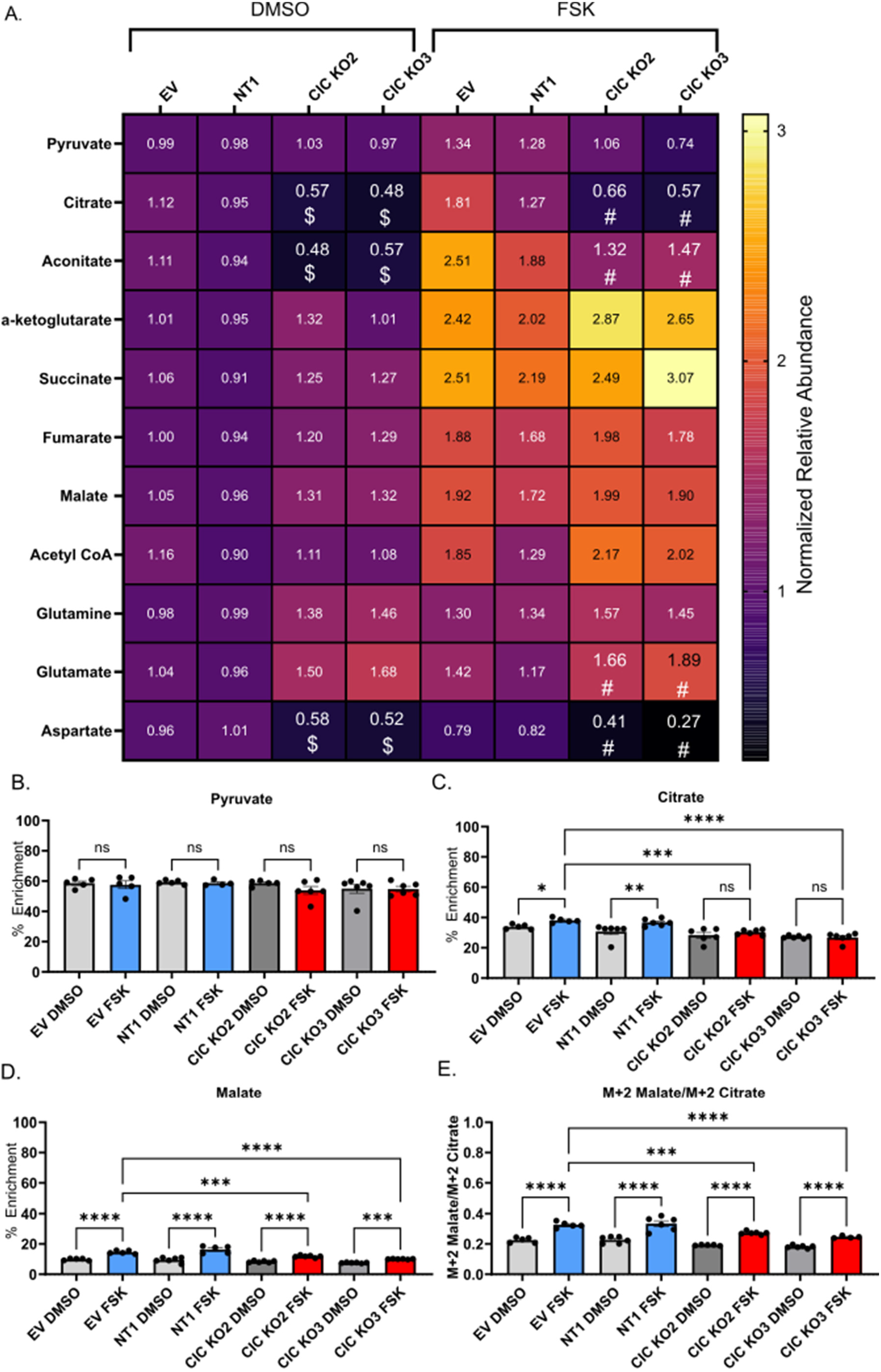
Loss of mitochondrial citrate efflux impairs metabolic reprogramming. A) Heat map of normalized relative metabolite abundance of DMSO and 40 µM forskolin treated empty-vector (EV), non-targeting, and CIC knockout BeWo cells. n=6 biologic replicates. Data are representative of mean +/- SEM. $, *p*<0.01 compared to EV-DMSO and #, p<0.01 compared to EV-FSK using ANOVA. B-D) Percent enrichment of [U-^13^C_6_]-glucose in B) Pyruvate C) Citrate and D) Malate following treatment of BeWo cells with DMSO or forskolin. E) Ratio of M+2 Malate/M+2 Citrate. n=6 biological replicates. Data are representative of mean +/-SEM. *, *p*<0.05; **, *p*<0.01; ***, *p*<0.001; and ****, *p*<0.0001.

Finally, to examine the impact of differentiation on flux of glucose-derived carbon into the second span of the TCA cycle, we examined the ratio of M+2 malate to M+2 citrate following loss of CIC. In control cells, the M+2 malate to M+2 citrate ratio increased following forskolin stimulation, suggesting increased incorporation of glucose-derived carbon into the second span of the TCA cycle **(Figure 7E)**. In CIC knockout cells, the M+2 malate to M+2 citrate ratio also increased, but to a smaller extent than observed in empty vector or scrambled gRNA-treated control cells. Combined, these findings demonstrate that loss of CIC contributes to metabolic adaptation and its absence decreases forward flux of glucose-derived citrate through the TCA cycle with differentiation.

## Discussion

Trophoblast differentiation requires coordinated metabolic, transcriptional and epigenetic reprogramming. Our results are the first to demonstrate that trophoblast differentiation is associated with changes in citrate metabolism and that loss of mitochondrial citrate export impairs trophoblast differentiation likely in part through dysregulation of histone acetylation. Overall, our work highlights specific metabolic differences between cytotrophoblasts and syncytiotrophoblasts and identifies the mitochondrial citrate carrier as a regulator of trophoblast differentiation.

Prior work highlights differences in cytotrophoblast and syncytiotrophoblast metabolism using Seahorse respirometry.^17,18^ Our work specifically demonstrates that differentiation is associated with increased ATP concentration and increased TCA cycle metabolite abundance and enrichment from ^13^C-labeled substrate. Based on our carbon tracing studies, cytotrophoblasts efflux mitochondrial citrate to a greater extent than syncytiotrophoblasts, whereas syncytiotrophoblasts preferentially retain citrate within the TCA cycle to a greater extent. This is consistent with recent reports highlighting a “non-canonical” TCA cycle in undifferentiated cells likely reflecting the activity of the citrate-malate shuttle necessary to sustain cytoplasmic acetyl-CoA pools in undifferentiated states to one supporting ATP generation following differentiation.^25^ Surprisingly, loss of mitochondrial citrate efflux decreased total citrate abundance, but not α-ketoglutarate or malate. Additionally, the M+2 malate to M+2 citrate ratio was not increased following loss of citrate efflux via CIC knockout. Together, this suggests that other anaplerotic sources such as glutamine may contribute to TCA cycle intermediates following loss of citrate efflux.

Acetyl-CoA cannot be transported across the mitochondrial inner membrane, thus citrate is a key metabolic intermediate though which carbon obtained from glucose metabolism contributes to cytoplasmic acetyl-CoA pools. Once in the cytoplasm, ACLY converts citrate to acetyl-CoA that can be used for histone acetylation.^34^ The increased citrate and acetyl-CoA with differentiation likely reflects an increase in mitochondrial metabolite pool size. Mitochondrial compartmentalization of citrate and acetyl-CoA may decrease substrate availability for histone acetylation. While our studies do not formally delineate metabolite location, defining the spatial-temporal regulation of acetyl-CoA pools involved in nuclear signaling events remains of interest for future investigation.^35,36^

While acetyl-CoA pool sizes are preserved in CIC knockout cells despite lower abundance of cellular citrate, it is unclear if this is due to increased acetyl-CoA compartmentalization or the result of other metabolic adaptations. Application of exogenous acetate partially rescues gene expression in CIC knockout cells, suggesting that cytoplasmic acetyl-CoA may be dysregulated in CIC knockout cells. In other systems, loss of ACLY is associated with increased expression of ACSS2, and increased ACSS2 activity cannot be excluded in CIC knockout cells.^30^ While loss of CIC impairs histone de-acetylation following differentiation, our work cannot delineate if this is the cause or result of impaired differentiation following loss of citrate efflux. Trophoblast differentiation is associated with decreased total histone acetylation but increased histone acetylation near loci associated with syncytiotrophoblast differentiation and decreased acetylation at loci associated with the less differentiated cytotrophoblast state.^11^ We observed global changes in histone acetylation following loss of mitochondrial citrate export and postulate that loss of citrate efflux impairs locus-specific regulation resulting in the observed impairments in trophoblast differentiation. Future work will investigate how loss of citrate efflux impacts locus specific regulation of gene expression during trophoblast differentiation.

This work focuses on how citrate may regulate gene expression through its action on acetyl-CoA pools and potentially histone acetylation. It is possible that changes in other metabolites such as α-ketoglutarate may impact histone methylation events and contribute to changes in gene regulation. α-ketoglutarate is a key cofactor required for demethylation by Jumanji demethylases and histone demethylation regulates trophoblast differentiation^37,38^. Additionally, the impact of other metabolic nodes such as glutamine, glutamate, fatty acids, and carnitines may facilitate adaptation to impaired mitochondrial citrate efflux.^36^ Future work will define how mitochondrial nutrient flux regulates histone acetylation and methylation events and whether changes in carbon sources specifically regulate this process.

While BeWo cells represent an established model to study syncytiotrophoblast differentiation and have strong genetic concordance with primary trophoblasts,^39^ the BeWo model may not fully recapitulate all aspects of trophoblast differentiation. However, similar decreases in *SLC25A1* were observed in trophoblast stem cells. Additionally, our finding of decreased histone acetylation upon differentiation is similar to prior work that demonstrates differentiation-dependent decreases in histone acetylation among primary trophoblasts, trophoblast stem cells and BeWo cells.^11^ Finally, our metabolic data aligns with work by others that suggests syncytiotrophoblasts are more metabolically active than cytotrophoblasts.^18^

Overall, this work provides evidence that mitochondrial citrate metabolism is regulated during differentiation and that differentiation is associated with decreased citrate efflux and increased forward flux through the TCA cycle. Loss of CIC results in impaired biochemical differentiation and broad changes in gene expression, though does not impair morphologic differentiation. Though the specific mechanisms of this gene dysregulation remain unclear, abnormalities in histone deacetylation may contribute to the broad gene expression differences. Collectively, this work highlights roles beyond ATP generation for mitochondria in trophoblasts and identifies mitochondrial citrate efflux as a key regulator of trophoblast differentiation.

## Acknowledgements

S.A.W. is supported by the Reproductive Scientist Development Program (RSDP) by the Eunice Kennedy Shriver National Institute of Child Health and Human Development (K12 HD000849) and the American Board of Obstetrics and Gynecology. P.A.C. is supported by R01DK091538 and R01AG069781. E.B.T. is supported by R01DK104998. M.D.G. is supported by 5R21HD102770. C.C.H. is supported by American Cancer Society Institutional Research Grant (IRG-21-049-61-IRG131). A.J.R. is supported by American Heart Association Career Development Award (CDA851976). This work was supported by the resources and staff at the University of Minnesota Genomics Center (https://genomics.umn.edu), Informatics Institute, and Imaging Centers (SCR_020997). The authors wish to thank Dr. Yoel Sadovsky, at Magee Women’s Research Institute, for helpful discussions throughout this project.

## Supplement Figure Legends

**Supplemental Figure 1:**
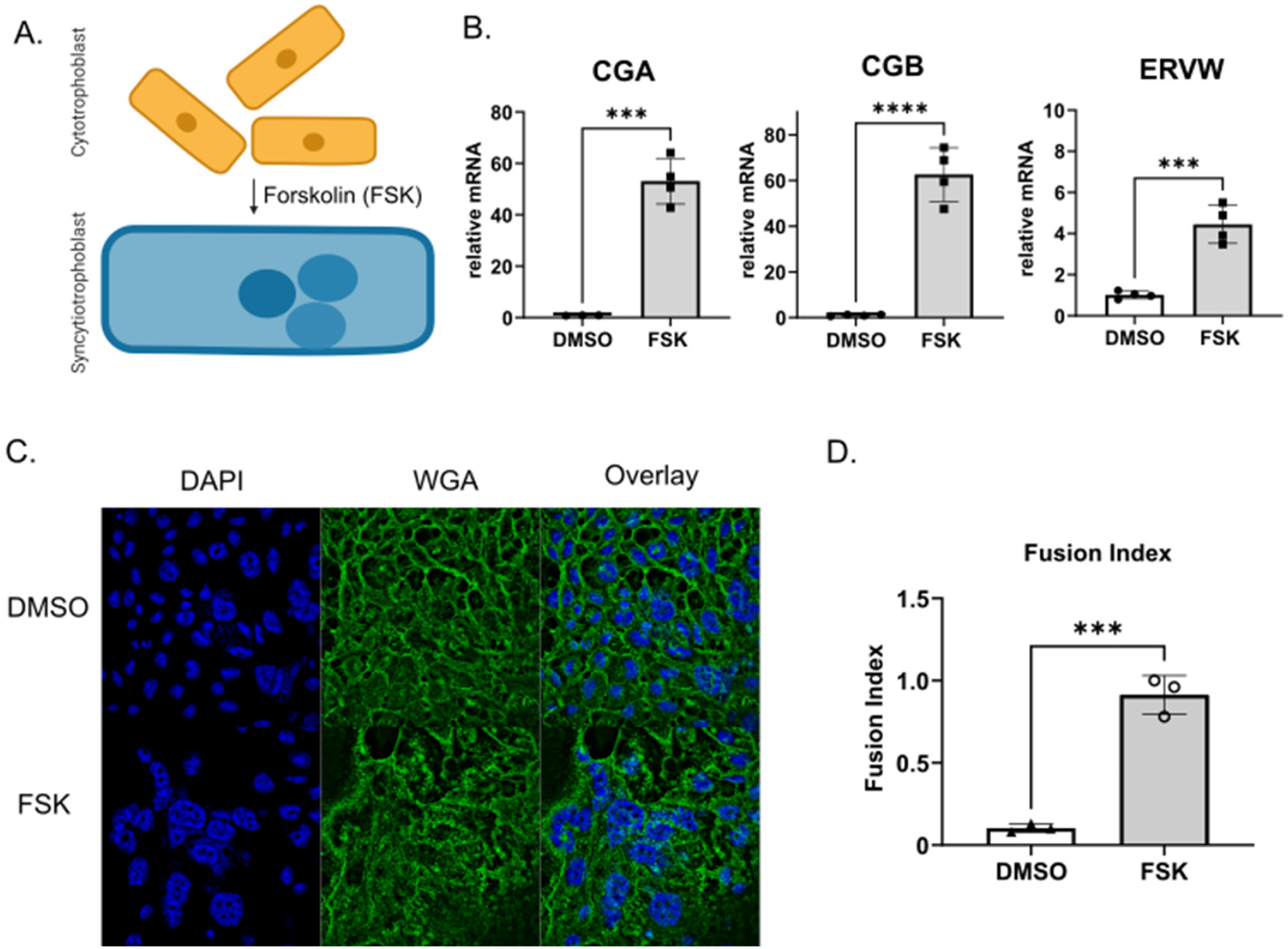
Confirmation of biochemical and morphologic differentiation In BeWo trophoblast cells. A) Schematic of cytotrophoblast fusion into multinucleated syncytiotrophoblasts (Figure created in Biorender) B) qPCR analysis showing upregulation of *CGA, CBG* and *ERVW* gene expression in response to differentiation, n=4; C) Representative images demonstrating morphologic changes associated with forskolin treatment. Blue depicts DAPI nuclear staining and green demonstrates wheat germ agglutinin staining of cell membrane. D) Quantification of Fusion Index (number of nuclei in syncytia/total number of nuclei). n=3. Data are representative of mean +/-SEM. ***, p<0.001; ****, p<0.0001.

**Supplemental Figure 2:**
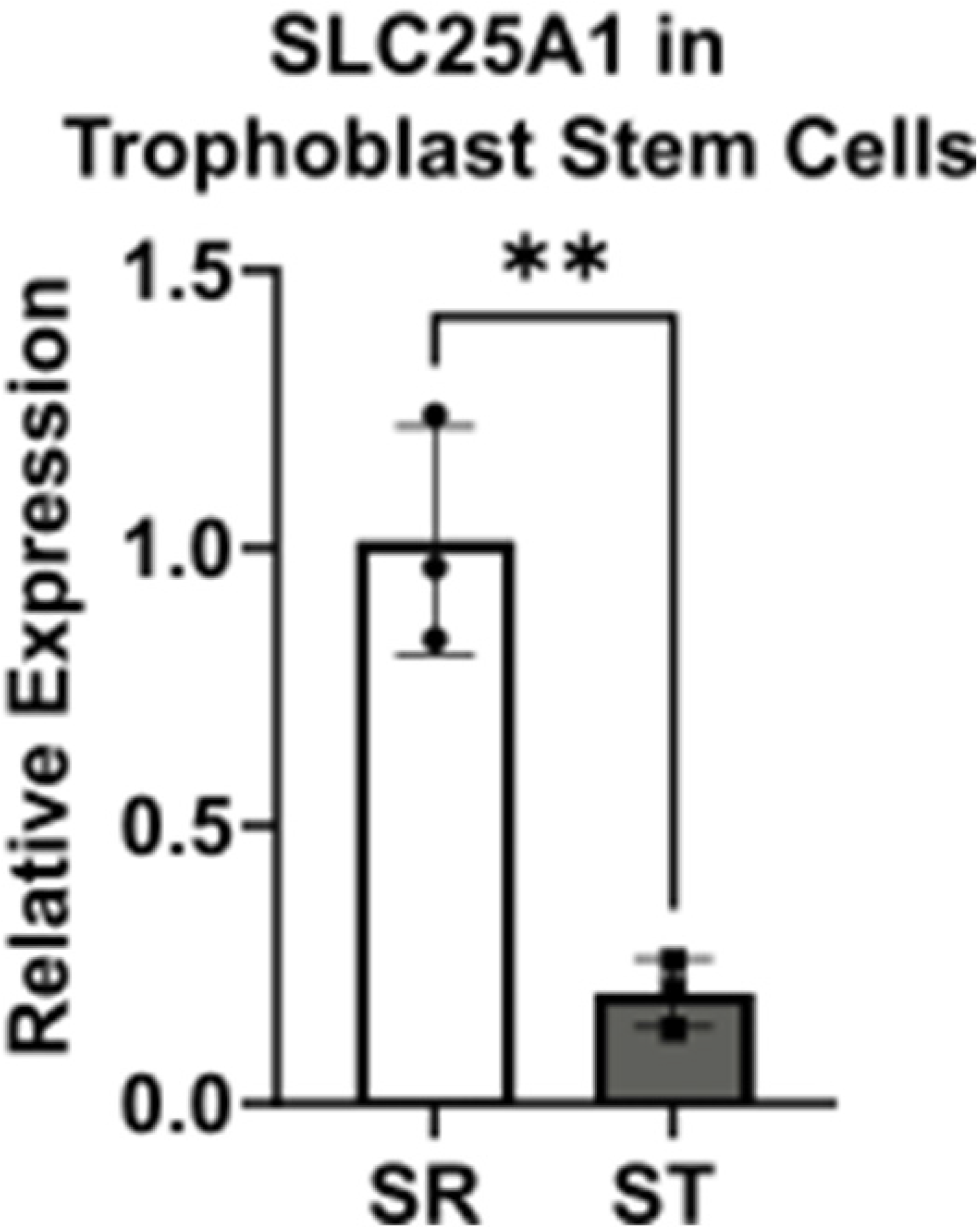
Decreased Expression of SLC25A1 in Trophoblast Stem Cell model of differentiation. A) qPCR demonstrating decreased *SLC25A1* expression following differentiation. SR=self renewing trophoblast stem cell. ST=syncytiotrophoblast. n=3. Data are representative of mean +/-SEM. **, *p*<0.01.

**Supplemental Figure 3:**
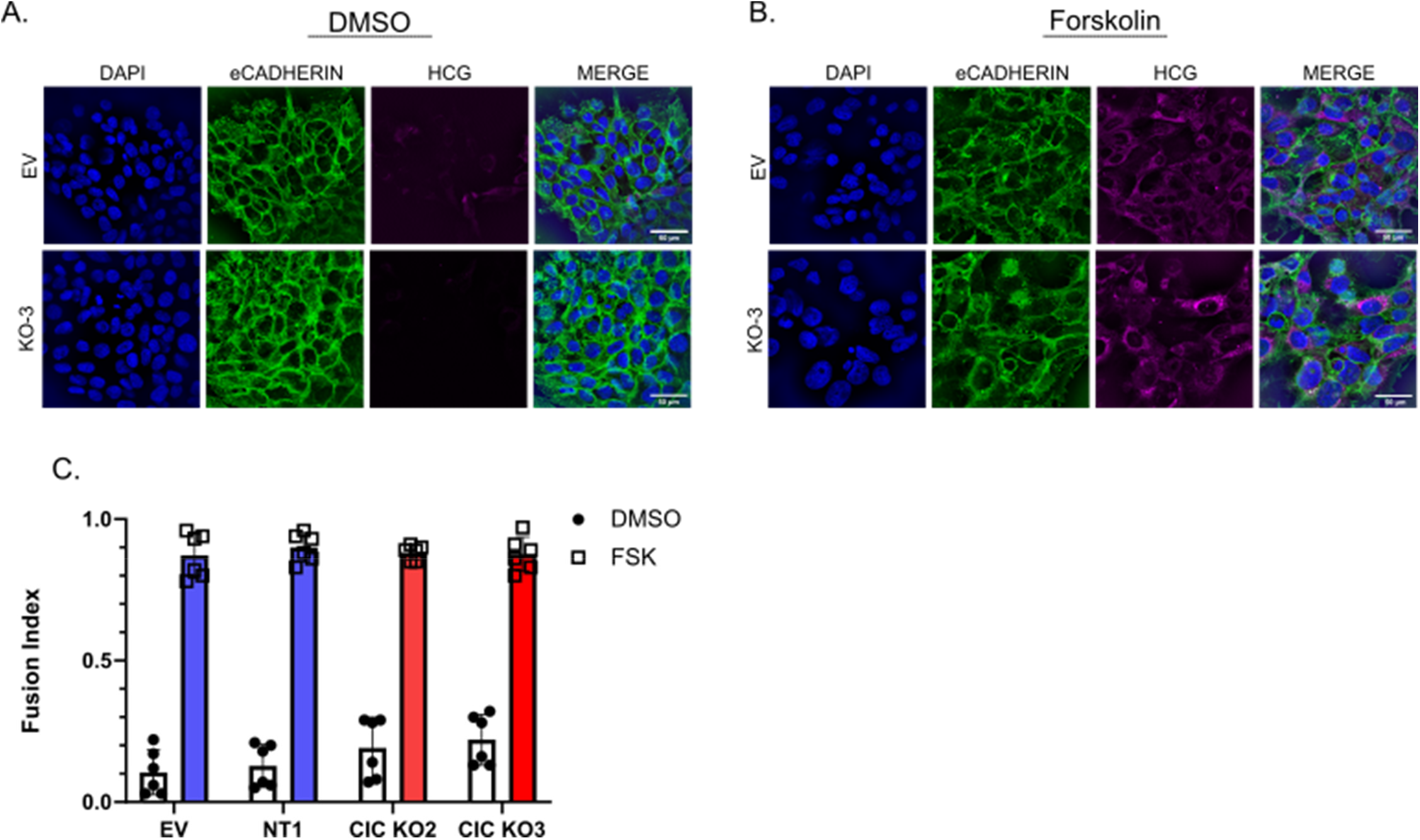
Loss of CIC does not impair morphologic syncytialization. Representative images demonstrating cell morphology following treatment of BeWo control or CIC Knockout cells with A) DMSO or B) 40 µM forskolin. Blue depicts DAPI nuclear staining, green demonstrates E-cadherin, and magenta represents HCG. Scale bar represents 50 µm. C) Quantification of Fusion Index (number of nuclei in syncytia/total number of nuclei). n=6.

## Methods

### STAR (Structure, Transparent, Accessible, Reporting) Methods

#### Key Resource Table

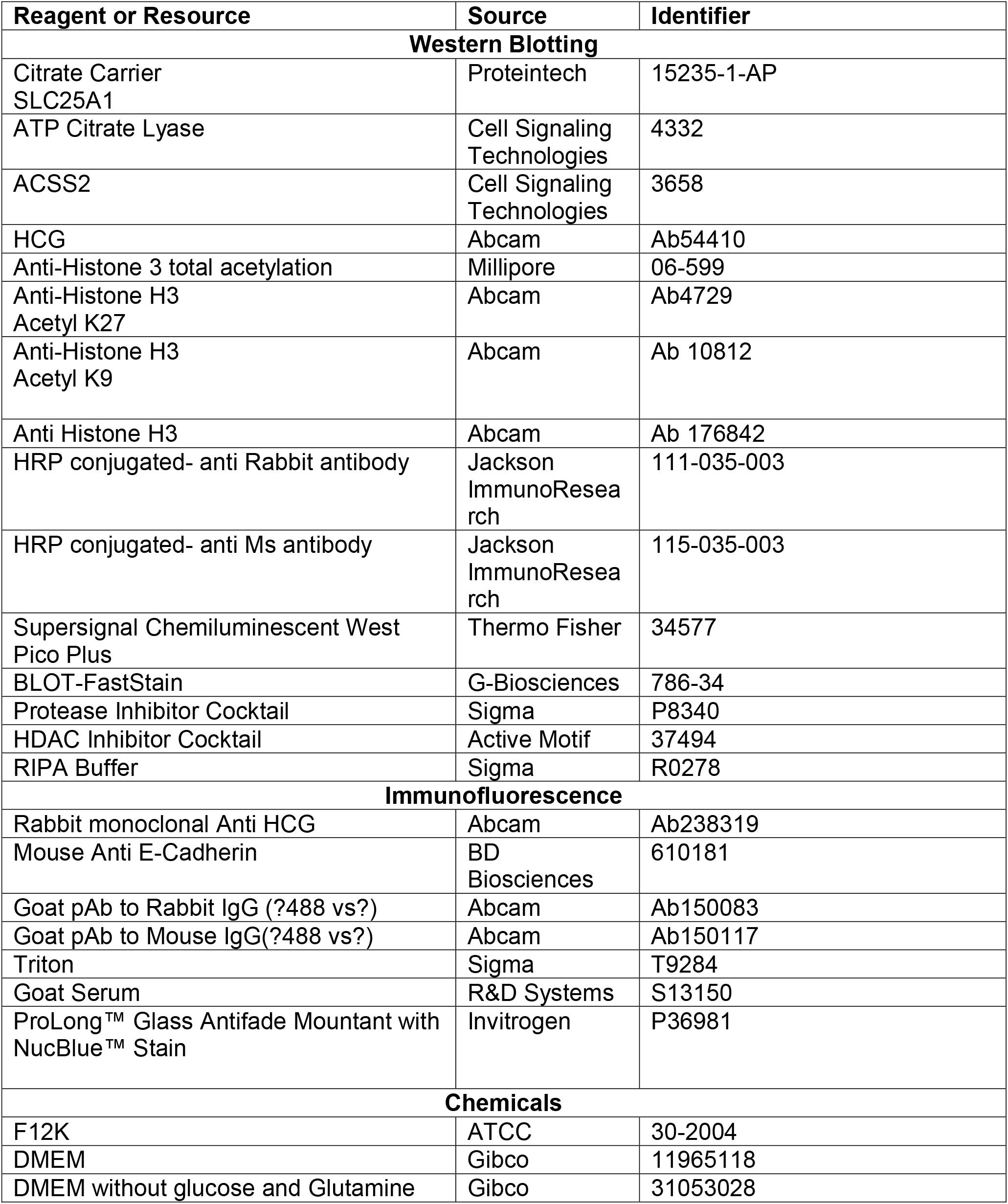
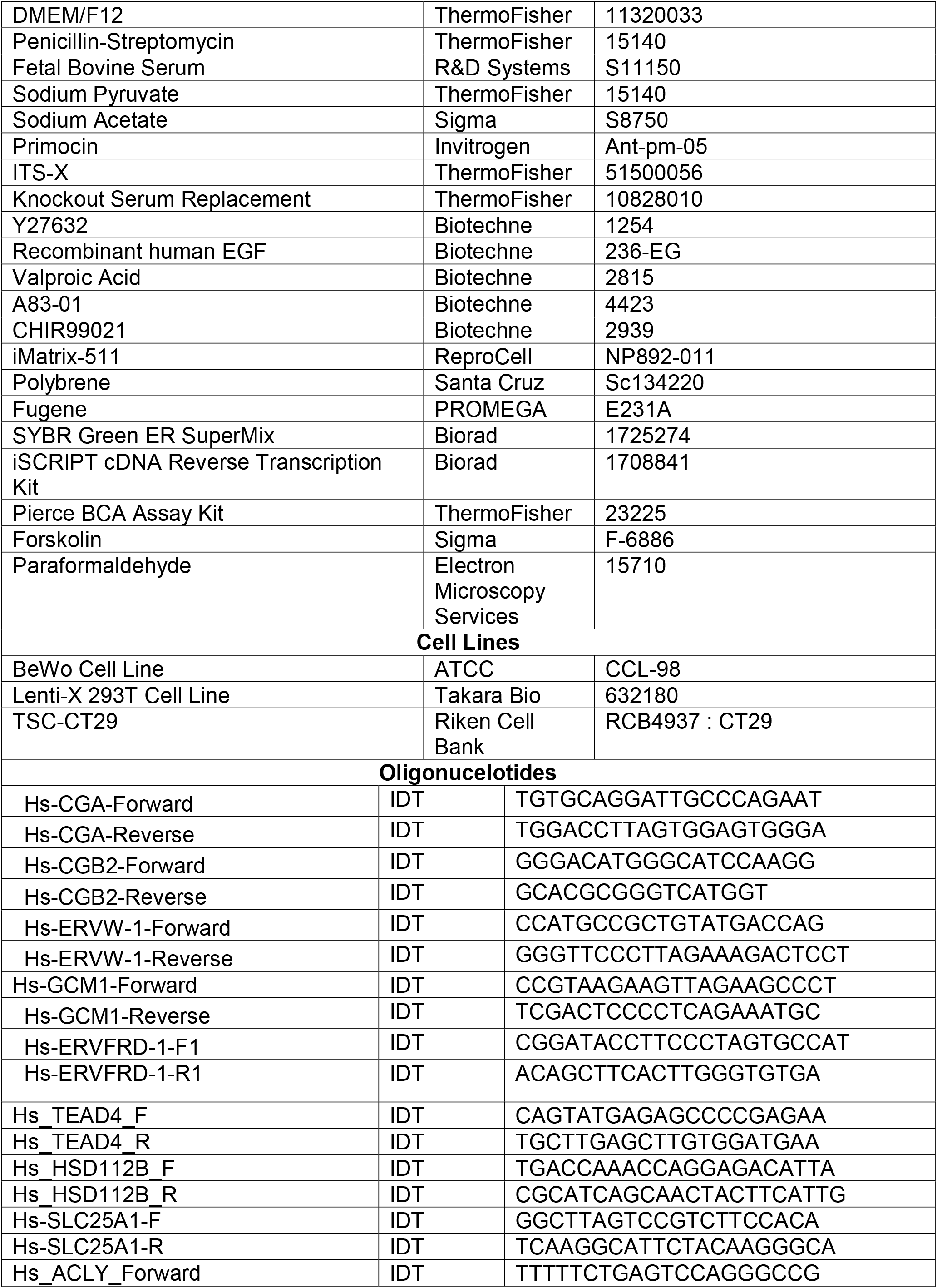
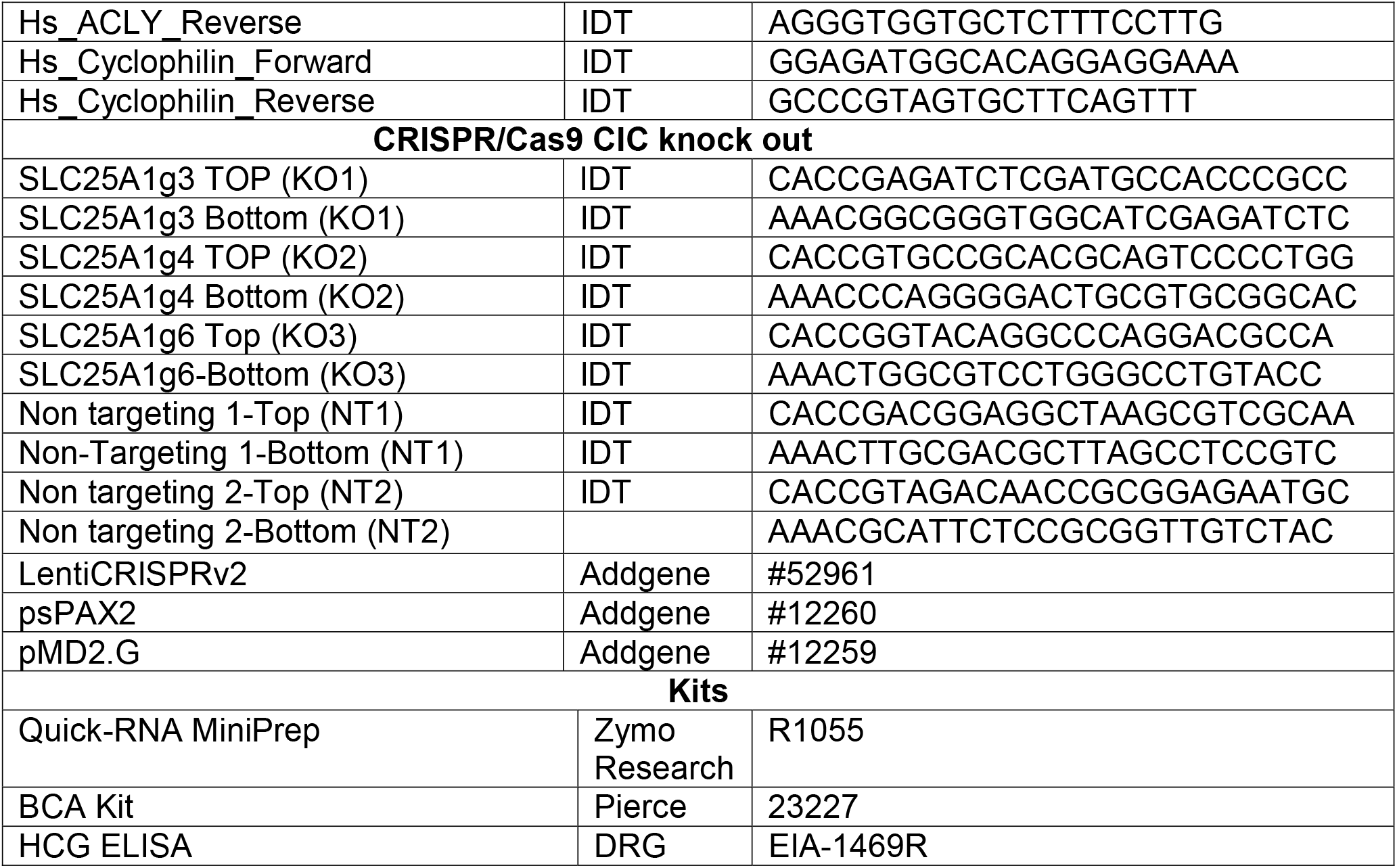

#### Lead Contact and Material Availability

Further information and requests for resources and reagents should be directed to Sarah Wernimont MD, PhD.

#### Experimental Model and Subject Details

No human subjects were used in this research.

#### Cell Lines

BeWo cells were purchased from ATCC. LentiX cells were purchased from Takara. TSC (CT-29 clone) were obtained from Rikken Cell Bank.

### Method Details

#### Cell Culture and Reagents

BeWo cells were purchased from ATCC and maintained in culture with equal parts F12K (ATCC) (supplemented with 10% Fetal Bovine Serum (FBS, Atlantic Biologics), Penicillin-Streptomycin (Life Technologies)) and DMEM Complete (4.5 g/L glucose, Life Technologies) (supplemented with glutamine (LifeTechnologies), sodium pyruvate (LifeTechnologies), 10% fetal bovine serum (Atlantic Biologics), and Penicillin-Streptomycin (Life Technologies)). To induce differentiation, BeWo cells were cultured for 24 hours on tissue culture treated plates and DMSO (vehicle control, 0.4%) or 40 uM forskolin (Sigma) was added. Media and treatment were replaced at 24 hours during differentiation and cells were harvested for analysis 48 hours after treatment. Lenti-X cells were purchased from Takara and maintained in DMEM containing 10% FBS, Pen/Strep and Sodium Pyruvate.

Trophoblast stem cells (TSC) were maintained in DMEM/F12 basal media containing with 100 µg/ml primocin, 0.15% BSA, 1% ITS-X, 1% Knockout Serum Replacement and 0.2 mM ascorbic acid and supplemented with 2.5 µM Y27632, 25 ng/ml EGF, 0.8 mM Valproic Acid, 5 µM A83-01, and 2 uM CHIR99021 on iMatrix-511 coated plates to maintain cells in self-renewing state. Cells were cultured at 37C, 5% CO2. To induce differentiation, 100,000 cells were plated on a 6 well plate coated with iMatrix-511 in self-renewing media. On day 1, media was replaced with DMEM/F12 basal media containing 100 µg/ml Primocin, 0.1% BSA, 1% ITS-X supplemented with 2.5 uM Y27632, 4% KSR, and 2 µM forskolin to induce differentiation. Self-renewing cells were maintained in self-renewing media. On day 3 following differentiation, RNA was isolated for qPCR.

#### Generation of SLC25A1 Knockout Cell lines

To generate CIC knockout BeWo cells, CRISPR guide RNAs (gRNAs) targeting exon 1 of *SLC25A1* were subcloned into lentiCRISPR v2 (Addgene Plasma 52961) using BbsI sites as described.^33,40^ Non-targeting guides were generated using previously reported sequences.

Lentivirus was made by transfecting LentiX Cells (Takara) with lentiCRISPRv2 vector including guide of interest, psPAX2 (Addgene plasmid #12260) and pMD2.G (Addgene plasma #12259) using Fugene transfection reagent (Promega) in DMEM containing 10% FBS and Pen/Strep. 24 hours after transfection, additional FBS was added to a final concentration of 30%. Virus containing media was harvested 72 hours after transfection, centrifuged to remove cellular debris and passed through a 0.45 uM filter (Pall Corporation). 1 ml of virus containing media was mixed with 1 ml of F12K and 1 µg/ml polybrene and added to BeWo cells plated 1 day prior to transduction. Forty eight hours after transduction, selection was started with 1 µg/ml puromycin and 1 mM acetate was added to media to support cell growth. Complete loss of CIC expression took approximately 12 days, likely due to slow turnover of mitochondrial transport proteins. After confirmation of CIC knockout, cells were used experimentally on days 14-28. Cells were maintained in culture in F12K:DMEM supplemented with 1 mM acetate. However, all differentiation experiments were done without supplemental acetate unless otherwise indicated.

#### Metabolomic Profiling and Isotope Tracing Metabolomics

##### Cell treatments and Harvesting

250,000 BeWo cells were plated and twenty four hours after plating, media was changed and cells were treated with 40 µM Forskolin or DMSO. Forty eight hours after initiation of treatment, cells were harvested for static metabolomics analysis. For isotope tracing metabolomics, cells were incubated with F12K and DMEM media supplemented with 17 mM [U-^13^C_6_]glucose (Cambridge Isotopes) or unlabeled naturally occurring glucose (Sigma) (standard concentration in F12K/DMEM media) for 6 hours. A mirror plate was maintained for each condition to facilitate normalization to total protein. After treatment, samples were collected by washing twice with ice cold PBS, once with ice cold water and snap freezing in liquid nitrogen prior to transferring cells in methanol to a fresh tube. Solvent was evaporated and samples stored at −80LC until ready for analysis.

Prior to analysis, metabolites were extracted by adding 1000 µL of 2:2:1 (v:v:v) Acetonitrile (AcN):Water(H_2_O):MeOH as previously described^41,42^. Samples underwent three rounds of vortexing, sonication and snap freezing in liquid nitrogen. The samples were incubated at −20°C for 1-4 hours, spun to remove proteins, transferred to fresh tube, evaporated, and reconstituted in 40 μL of 1:1 (v:v) AcN:H_2_O. Prepared samples were vortexed, sonicated, centrifuged, and analyzed. All analyses were performed on Thermo Vanquish liquid chromatograph hyphenated with Thermo Q-Exactive Plus mass spectrometer equipped with heated ESI (HESI) source.

##### Relative Abundance of glycolytic and TCA cycle intermediates

To identify selected TCA cycle and glycolytic metabolites, samples were analyzed using Atlantis Premier BEH Z-HILIC Column (2.1 mm × 100 mm, 1,7 μm). Separation was complete using Mobile phase A (15 mM ammonium bicarbonate in water, pH 9.0) and Mobile Phase B (15 mM ammonium bicarbonate pH, 9.0 with 90% AcN) using the following gradients: 10% Mobile Phase A for 5 min, 35% Mobile Phase A for 2 min and 10% Mobile Phase A for 3 minutes. Separations were performed at a flow rate 0.5 to 1 mL/min, column temperature at 30°C and an injection volume of 2 μL. The mass spectrometer operated in negative mode using full scan (FS) mode (*m/z* 68–1,020) with optimized HESI source conditions: auxiliary gas 10, sweep gas 1, sheath gas flow at 30 (arbitrary unit), spray voltage −4 kV, capillary temperature 350°C, S-lens RF 50, and auxiliary gas temperature 350°C. The automatic gain control (AGC) target was set at 3e6 ions and resolution was 70,000.

Xcalibur Quan Browser was used for peak identification and integration. Metabolite profiling data was analyzed with QuanBrowser using standard verified peaks and retention times. Metabolites peak identity was confirmed based on retention time, *m/z* and compared to authentic internal standards as previously described.^42,43^ Integrated signal for each metabolite was normalized to total protein determined using Bicinchoninic Acid Assay (BCA) on mirror plates for each condition. All normalized signal intensity was normalized to DMSO control conditions and differences in relative abundance determined.

##### Isotope Tracing Untargeted Metabolomics

For isotope tracing experiments, following metabolite extraction, 20 mM of ammonium phosphate was added to each sample and samples were analyzed using on a SeQuant ZIC-pHILIC (Merc) column (150 mm X 2.1 mm) as previously described.^42-44^ Briefly, separation was performed using binary gradients of mobile phase A (10 mM Ammonium acetate/ammonium hydroxide, pH 9.0, 95% water/5% AcN) and mobile phase B (AcN). Following gradient was used: 100% to 0% B for 50 minutes, 0% B for 7 minutes and 100% B for 13 minutes. Flow rate was set to 0.2 mL/min, column temperature was 45LC, and injection volume was 2 µL. The mass spectrometer operated in negative mode using full scan (FS) mode (*m/z* 68–1,020) using optimized HESI source conditions: auxiliary gas 10, sweep gas 1, sheath gas flow at 30 (arbitrary unit), spray voltage −4 kV, capillary temperature 350°C, S-lens RF 50, and auxiliary gas temperature 350°C. The automatic gain control (AGC) target was set at 3e6 ions and resolution was 70,000.

Compound Discoverer 3.1(Thermo) was used for peak identification, integration and natural abundance correction. Briefly, negative mode data processing used the following parameters: 5 ppm mass tolerance, 1 minute maximum retention drift time, minimum scans per peak of 5 and maximum peak width of 0.5 minutes. Background signals were excluded based on the Sample/Blank signal ratio >3. Percent (%) enrichment was determined for each metabolite of interest. For poorly integrated peaks, Xcalibur was used for isotopologue peak identification and integration following natural abundance correction by IsoCorrectoR.^45^

##### Energy Metabolite Quantification

To quantify energy metabolites using methods described in^46,47^, cell pellets were treated with 0.4 M perchloric acid, 0.5 mM EGTA extraction solution containing [^13^C_10_, ^15^N_5_]ATP sodium salt (100 μM), [^13^C_10_,^15^N _5_]AMP sodium salt (100 μM), and [1,2-^13^C_2_]acetyl-CoA lithium salt (5 μM)) as internal standards, all purchased from Sigma. Samples underwent three rounds of vortexing, snap freezing in liquid nitrogen, and sonication. Following 10 minute incubation on ice, samples were centrifuged and supernatants was neutralized with 0.5 M K_2_CO_3_. Following additional centrifugation, extracts were analyzed by LC-MS/MS as previously described and normalized to total protein for each condition.^46,47^

#### Quantitative PCR

Total RNA was extracted from cells using the Quick-RNA MiniPrep protocol provided by the kit’s manufacturer (Zymo) and 1 µg of RNA was used to synthesize cDNA (iSCRIPT, BioRad). For quantitative real time PCR, reactions were carried out using SsoAdvanced Universal SYBR Green Supermix (BioRad) on a CFX384 Real-Time System (Bio-Rad). Transcripts were quantified using the 2^-^ΔΔ_CT method._ and normalized to cyclophiln. Primer sequences are listed in Key Resources.

#### RNA-Sequencing

RNA was isolated from control and CIC knock out cell lines following treatment with DMSO or 40 uM forskolin for 48 hours using Quick-RNA MiniPrep (Zymo). The University of Minnesota Genomics Core assessed sample quality using Agilent BioAnalyzer 2100 and all samples were found to have an RNA Integrity Number (RIN) of 8 or greater. Libraries were created using Illimunia TruSeq RNA sample preparation (Cat. # RS-122-2101) kit following manufacturer’s instructions. Pool libraries were loaded onto a NovaSeq flow cell and sequenced. Base call files for each cycle of sequencing were generated by Ilumina Real Time Analysis Software and stored on servers at the Minnesota Super Computing Institute. Primary analysis and de-multiplexing was performed using Illumina’s bcl2gastq v2.20. 2×150bp FastQ paired-end reads for 12 samples (n=54.6 Million average per sample) were trimmed using Trimmomatic (v0.33) enabled with the optional “-q” option; 3bp sliding-window trimming from 3’ end requiring minimum Q30. Quality control was completed by FastQC. Read mapping was performed via Hisat2 (v2.1.0) using the human genome (GRCh38) as reference. Gene quantification of Ensembl v97 annotations was completed using Subread. Counts were normalized using the median ratio method and differentially expressed genes were identified using the Wald test in DeSeq2.^48^ The Benjamini and Hochberg method was used to account for False Discovery Rate in DeSeq2 and give an adjusted *p*. Genes with an adjusted *p*-value ≤ 0.01 were considered statistically significant. PCA plots were made using ggplot2^49^, volcano plots made using EnchanedVolcano^50^, and Pathway analysis completed using cluterProfiler ^51^ in R(v4.2.1). RNA Seq Data has been deposited at NCBI.

#### Protein Expression

Following treatment, cells were washed with ice cold phosphate buffered saline (PBS) and lysed using 250 ul of RIPA Buffer (Sigma) supplemented with 1X Protease Arrest inhibitor. For assessment of histone acetylation, 1X HDAC inhibitor (Active Motif) was added to lysis buffer, and cells were sonicated for 45 seconds with 3 second pulses at power of 30% on ice. Lysates were centrifuged for 5 min at 4°C at 21,000x*g*. Bicinchoninic Acid Assay (Sigma) was used to determine protein concentration. 5 µg of protein was used for histone expression and 30 µg of protein was used to detect expression of other proteins. Lysates were run on 4-12% or 10% Bis-Acrylamine precast gels (Invitrogen) and transferred to nitrocellulose. Membranes were blocked with 2% bovine serum albumin (Sigma) or 5% milk in tris buffered saline (TBS) and stained overnight at 4°C with primary antibodies (see key resources). Blots were washed with TBS and stained with HRP-tagged secondary antibodies. Blots were developed using SuperSignal West Pico PLUS Chemiluminescent Substrate (Thermo). Total protein loading was assessed by BlotFastStain (G-Biosciences). Blots were imaged using Bio-Rad CheiDoc MP imaging system and band density was quantified using ImageJ.

#### Imaging

50,000 BeWo cells were plated on poly-L-lysine treated glass coverslips coated with 10% FBS in a 24-well plate. At 24 hours, media was replaced and cells were treated with 40 µM forskolin or DMSO as vehicle control. At 48 hours, wells were washed with PBS and were fixed with 4% paraformaldehyde (Electron Microscopy Services) for 10 minutes at 37°C. Wells were washed again twice with PBS, then each coverslip was moved to a staining box. The cells were permeabilized with 0.1% TritonX (Sigma) and incubated for 10 minutes at room temperature. The coverslips were again washed twice with PBS, then incubated with 5% goat serum at room temperature for an hour. After one hour, coverslips were washed twice with PBS then incubated with mouse E-cadherin (BD Biosciences) and rabbit monoclonal HCG (Abcam) primary antibodies (Abcam) (diluted 1:200) overnight at 4°C protected from light. The following day, the coverslips were washed three times with PBS and incubated with mouse and rabbit secondary antibodies (Abcam) (diluted 1:400) for 1 hour, protected from light at room temperature. Coverslips were washed with PBS, dipped in MilliQ water, and placed on slides with Prolong Glass Antifade Mounting Media with DAPI (Invitrogen). Slides were allowed to dry overnight and then sealed using clear nail polish. Coverslips were kept at –20°C until imaging. Images were acquired on an Olympus FluoView BX2 Upright Confocal at the University of Minnesota Imaging Core. Fusion index was quantified as the # of nuclei in syncytia divided by the total # of nuclei using ImageJ.

#### HCG ELISA

250,000 BeWo cells were plated and treated with DMSO or 40 uM forskolin to induce syncytialization. Forty eight hours after treatment, media samples were collected and ELISA completed following manufacturer’s instructions.

#### Quantification and Statistical Analysis

The Student’s *t* test was used when comparing the means of two groups after a limited number of statistical outliers were excluded following identification by the Grubbs test. ANOVA was used to compare the means of more than two groups. *p* values less than 0.05 were considered statistically significant. Data were organized and analyzed using Microsoft Excel and GraphPad Prism.

